# A *Saccharomyces cerevisiae* Model for Overexpression of Ntg1 a Base Excision DNA Repair Protein Reveals Novel Genetic Interactions

**DOI:** 10.1101/2022.05.06.490893

**Authors:** Annie J. McPherson, Ziad M. Jowhar, Paul W. Doetsch, Anita H. Corbett

**Affiliations:** Department of Biology, Emory University, Atlanta, GA, 30322, USA; Genetics and Molecular Biology Graduate Program, Emory University, Atlanta, GA, 30322, USA; Initiative for Maximizing Student Development, Emory University, Atlanta, GA, 30322, USA; Mutagenesis and DNA Repair Regulation Group, National Institutes of Environmental Health Sciences, Division of Intramural Research, Genome Integrity & Structural Biology Laboratory, Durham, NC, 27709, USA

**Keywords:** Base excision repair, Ntg1, NTHL1, *S. cerevisiae*, sumoylation

## Abstract

The Base Excision Repair (BER) pathway repairs oxidative DNA damage, a common and detrimental form of damage to the genome. Although biochemical steps in BER have been well define, little is understood about how the pathway is regulated. Such regulation is critical, as cells must respond rapidly to DNA damage while avoiding aberrant activation of repair proteins that can produce DNA damage as intermediates in the repair pathway. Indeed, overexpression of the human BER protein, NTHL1, a DNA N-glycosylase, can cause genomic instability and early cellular hallmarks of cancer. We developed a *Saccharomyces cerevisiae* model to explore how overexpression of NTHL1 may impair cellular function. Overexpression of Ntg1, the budding yeast orthologue of NTHL1, impairs cell growth. To dissect mechanisms underlying this growth defect, we overexpressed either wild-type Ntg1 or a catalytically inactive variant of Ntg1 (ntg1_catdead_). Consistent with results obtained for NTHL1, both variants of Ntg1 impair cell growth, but only the wild-type protein causes accumulation of double-strand breaks and chromosome loss. We screened a panel of DNA repair mutants for resistance/sensitivity to overexpression of wild-type Ntg1 or ntg1_catdead_. This analysis identified several cellular pathways that protect cells from Ntg1-induced damage, providing insight into interplay between DNA repair pathways. Finally, we identified a link to sumoylation and probed how this post-translational modification could contribute to regulation of Ntg1 function. This study describes a budding yeast system to understand how cells regulate and respond to dysregulation of the BER pathway.

**Take Away:**

- Overexpression of a base excision DNA repair protein impairs cell growth
- Overexpression of a base excision DNA repair protein can cause DNA damage
- Multiple mechanisms cause DNA damage from overexpression of a repair protein
- DNA repair pathways functionally interact to protect cells from DNA damage
- Precise regulation of the activity of DNA repair proteins is critical

## INTRODUCTION

Within both the nucleus and the mitochondria, DNA is susceptible to damage (Altieri, Grillo, Maceroni, & Chichiarelli, 2008; Haag-Liautard et al., 2008; Hoeijmakers, 2009; Maynard, Schurman, Harboe, de Souza-Pinto, & Bohr, 2009; Miquel, 1991; Richter, Park, & Amest, 1988). There are many sources of both exogenous and endogenous DNA damage (Altieri et al., 2008; Chatterjee & Walker, 2017; Maynard et al., 2009; Mullenders, 2018; Turrens, 2003). An abundant endogenous source of DNA damage is reactive oxygen species (ROS) which are a byproduct of energy production within the cell (Altieri et al., 2008; Bauer, Corbett, & Doetsch, 2015; Finkel & Holbrook, 2000; Limón-Pacheco & Gonsebatt, 2009; Tomas Lindahl, 1993; Maynard et al., 2009; Riley, 1994). ROS can cause oxidative damage throughout the cell and can induce many types of DNA damage including 8-oxo-2’deoxyguanosine (Bauer et al., 2015; Beckman & Ames, 1997; Fraga, Shigenaga, Park, Degant, & Amest, 1990; Maynard et al., 2009). These damages must be rapidly and efficiently repaired to maintain genomic and genetic stability (Bauer et al., 2015; Boesch et al., 2011; Jeggo, Pearl, & Carr, 2016; Wei, 1998). To combat the myriad of damages that occur, cells employ a battery of DNA repair pathways (Altieri et al., 2008; Bauer et al., 2015; Boesch et al., 2011; Bohr, 2002; Chatterjee & Walker, 2017; Y. J. Kim & Wilson, 2012; Maynard et al., 2009; O’Rourke, Doudican, Mackereth, Doetsch, & Shadel, 2002).

Base excision repair (BER) is the major repair pathway for oxidative DNA damage (Bauer et al., 2015; Boesch et al., 2011; Y.-J. a. Kim, 2012; Maynard et al., 2009; Wallace, 2014). BER is initiated by an *N*-glycosylase detecting a non-helix distorting base damage (Y.-J. a. Kim, 2012; Tomas Lindahl, 1976; T. Lindahl, 1979). The *N*-glycosylase flips out the base and cleaves the base from the backbone, resulting in an abasic site^25,28,29^. The DNA backbone on the 5’-side of the abasic site is then cleaved by an apurinic/apyrimidinic (AP) endonuclease or on the 3’-side by a *N*-glycosylase with AP lyase function (Boiteux & Guillet, 2004; Y.-J. a. Kim, 2012). If cleavage occurs on the 3’-side of the abasic site, an AP endonuclease must cleave the backbone and expose the preceding hydroxyl, to allow DNA synthesis to occur (Boiteux & Guillet, 2004; Y.-J. a. Kim, 2012). After synthesis, the strand is ligated back together to complete the repair process (Bauer et al., 2015; Y.-J. a. Kim, 2012).

The BER proteins must be available for rapid deployment in response to DNA damage but must also be precisely regulated to prevent promiscuous damage to the genome (Christmann & Kaina, 2019; Knudsen, Andersen, Ltzen, Nielsen, & Rasmussen, 2009; K. L. Limpose, Corbett, & Doetsch, 2017; Saha, Rih, Roy, Ballal, & Rosen, 2010). One major protein that initiates DNA repair in the BER pathway is the *N*-glycoslyase NTHL1, an evolutionarily conserved member of the endonuclease III family (Aspinwall et al., 1997; Bandaru, Sunkara, Wallace, & Bond, 2002; Imai et al., 1998). NTHL1 is a bifunctional DNA *N*-glycosylase with associated apurinic/apyrimidinic (AP) lyase function (Ikeda et al., 1998; Marenstein et al., 2003). As NTHL1 initiates DNA repair by introducing a nick in the DNA backbone, its activity must be tightly regulated to prevent the accumulation of spurious nicks in the genomic material (K. L. Limpose et al., 2017).

As with a number of other DNA repair pathway proteins, mutations in the *NTHL1* gene have been linked to cancer (Larouche & Akirov, 2019; Terradas & Munoz-Torres, 2019; Tubbs & Nussenzweig, 2017; Weren et al., 2015). In addition, consistent with the concept that precise regulation of NTHL1 function is critical, recent work has revealed that NTHL1 expression is misregulated in a variety of types of cancer (Albertson, 2006; Goto et al., 2009; Koketsu, Watanabe, & Nagawa, 2004). Furthermore, a recent study demonstrated that overexpression of NTHL1 in non-cancerous human bronchial epithelial cells results in loss of genetic information and early hallmarks of cancer including genomic instability (Kristin L. Limpose et al., 2018). Interestingly, overexpression of a catalytically inactive form of NTHL1 induced similar phenotypes as wild-type NTHL1, suggesting multiple modes by which overexpression of this BER protein can induce DNA damage/genome instability. Overexpression of NTHL1 in this cell model triggered replication stress signaling. Studies using a reporter system revealed a decrease in homologous recombination in these cells overexpressing NTHL1. This work provided important insight into how overexpression of a BER protein could contribute to genomic instability but could not broadly define the spectrum of pathways cells may employ to respond to such damage (Kristin L. Limpose et al., 2018).

In the present study, we sought to develop budding yeast as a model to investigate how cells respond to damage induced by dysregulation of early steps in BER. To do so, we first established that overexpression of *S. cerevisiae* Ntg1, which is the budding yeast orthologue of NTHL1 (Eide et al., 1996), causes a growth defect when overexpressed in *S. cerevisiae*. Consistent with the studies of NTHL1 (Kristin L. Limpose et al., 2018), overexpression of catalytically inactive Ntg1 (ntg1_catdead_) also impairs cell growth. While overexpression of wild-type Ntg1 causes an increase in double strand breaks and chromosome loss, overexpression of catalytically inactive ntg1_catdead_ does not. These results suggest that the growth defects seen in cells that overexpress Ntg1 or ntg1_catdead_ are triggered, at least in part, by separate mechanisms. We then took advantage of the budding yeast system to probe genetic interactions with BER by screening a panel from the gene deletion collection for deletion mutants that show either enhanced sensitivity or resistance to overexpression of Ntg1 and/or ntg1_catdead_. We identified a number of pathways that show such genetic interactions, including the homologous recombination pathway, nucleotide excision repair, and SUMO-mediated DNA damage response. As previous work demonstrated that Ntg1 is sumoylated (Daniel B. Swartzlander et al., 2016), we expanded our analysis of the sumoylation pathway and also explored whether SUMO modification of Ntg1 impacts overexpression of Ntg1. By taking advantage of these genetic approaches, we have identified pathways of interest that could be explored to better understand the mechanism by which overexpression of human NTHL1 could contribute to cancer phenotypes.

## MATERIALS and METHODS

### Strains, Plasmids, and Media

All haploid *S. cerevisiae* strains and plasmids used in this study are listed in Table 1. *S. cerevisiae* cells were cultured at 25°C, 30°C, or 37°C in YPD medium (1% yeast extract, 2% peptone, 2% dextrose, 0.005% adenine sulfate, and 2% agar for plates) or SD medium (0.17% yeast nitrogen base, 0.5% ammonium sulfate, 0.5% adenine sulfate, and 2% agar for plates) or SD medium (0.17% yeast nitrogen base, 0.5% ammonium sulfate, 0.005% adenine sulfate, and 2% agar for plates) with either 2% dextrose, 3% raffinose, 2% galactose, 3% glycerol, or both 3% raffinose and 2% galactose. To introduce plasmids, cells were transformed by a modified lithium acetate method (Ito, Fukuda, Murata, & Kimura, 1983).

A galactose inducible 2μ vector (*pGAL1, URA3*), pPS293 (Addgene plasmid #8851; http://n2t.net/addgene:8851; RRID:Addgene 8851) was employed to express C-terminally epitope-tagged Ntg1-2xmyc (pAC3425). We expressed wild-type Ntg1, a catalytically inactive Ntg1 (ntg1_catdead_; *ntg1*_*K243Q*_) (Augeri, Hamilton, Martin, Yohannes, & Doetsch, 1994; D. B. Swartzlander et al., 2010; Daniel B. Swartzlander et al., 2016), a nonsumoylatable Ntg1 (ntg1ΔSUMO; *ntg1*_*K20,38,376,388,396R*_) (Daniel B. Swartzlander et al., 2016). This Ntg1 variant was also generated in the catalytically inactive form ntg1ΔSUMO_catdead_ (*ntg1*_*K20,38,376,388,396R,K243Q*_). The resulting plasmids were sequenced to ensure the correct desired sequence and the absence of any additional mutations.

The *S. cerevisiae* strain expressing endogenous Rad52-YFP (de Mayolo et al., 2010) was provided by Rodney Rothstein. This strain was employed to assay double strand break formation. The *S. cerevisiae* reporter strain containing a chromosome fragment with *SUP11* (Au, Crisp, Deluca, Rando, & Basrai, 2008) was provided by Munira Basrai. This strain was employed to quantify chromosome loss in response to overexpression of Ntg1. The *S. cerevisiae* haploid deletion collection was utilized to explore the genetic interactions with overexpression of Ntg1. The SUMO pathway mutant collection (E3 ligase mutant strains, *siz1Δ, siz2Δ*, and *siz1Δ/siz2Δ*, and desumoylase mutant stains *ulp1-1* and *ulp2Δ*) were utilized to assess the impact of global sumoylation loss or accumulation (Erica S. Johnson, 2004; E. S. Johnson & Gupta, 2001; Li & Hochstrasser, 1999).

### *S. cerevisiae* Growth Assays

*S. cerevisiae* cells containing *URA3* plasmids were grown in media lacking uracil plus 3% raffinose overnight. The next day, the OD600 of the cultures was measured and the cultures were diluted down to the lowest OD600. The cultures were then serially diluted 5-fold and 2.5 μL of each dilution was spotted onto plates lacking uracil containing either 2% glucose or 2% galactose. Plates were grown at 30°C and pictures were taken of the plates on day 2.

Liquid growth curves were collected for three independently isolated colonies per sample. Cells were grown overnight at 30°C in media lacking uracil with 3% raffinose. Cell concentrations were normalized by OD600, and then samples were diluted to an OD600 of 0.05 in 150 µL of media lacking uracil with either 2% glucose or 2% galactose and placed in the wells of a 96-well microtiter plate. Cell samples were loaded in triplicate, were grown at 30°C with shaking, and absorbance at OD600 was measured every 30 minutes for 24 hours in an ELX808 Ultra microplate reader with KCjunior software (Bio-Tek Instruments, Inc.).

### Cell Viability Assay

Cells were grown overnight in media lacking uracil plus 3% raffinose. The next day, cultures were harvested at mid-log phase (OD600 of 0.3-0.6) and equalized, then diluted 1:10 into media lacking uracil plus 2% galactose. Cells were grown in galactose overnight at 30°C. The next morning, OD600 was measured, cells were equalized again, diluted 1:1000 and an estimated 100 cells were plated onto plates lacking uracil plus 2% glucose and the plates were incubated for five days at 30°C. On day five, colonies were counted. The number of colonies grown on the plates containing Vector alone was considered the total number of live cells plated and therefore considered 100% survival. We then divided the number of colonies grown per plasmid by the total number of live cells plated and converted to a percentage. These experiments were conducted in both biological and technical triplicate. Standard deviations in the biological replicates are shown.

### Quantification of Cell Growth

Cells were grown, spotted, and imaged as described above. In order to better compare differences in growth, each strain and corresponding plasmid is given a value between 1 and 10 based on growth. A value of 10 corresponds to the growth of the control Vector for each mutant. A spot with large full colonies is given the value of 2 points, while spots with smaller colonies are given the value of 1. Spots with no growth are given a value of 0. This scoring approach is used for each of the five spots of serially diluted cells, yielding the range of 1-10 for each sample analyzed as we only analyzed samples where some growth could be detected in the most concentrated spot. These values, which are averages of the value obtained for biological triplicates, are displayed as a heat map.

### Double Strand Break Formation Assay

Cells containing an endogenous Rad52-YFP (de Mayolo et al., 2010) (generously provided by Rodney Rothstein) were grown overnight in media lacking uracil plus 3% raffinose. The next day, cells were diluted and grown until mid-log phase (OD600 0.3-0.6). At mid-log phase, the cells were dosed with 2% galactose and control samples with Vector alone were incubated in the presence of 2% galactose and 0.3% MMS. These cultures were incubated at 30°C for 2 hours. 30 minutes before treatment ended, DAPI was added to the culture to allow visualization of the nucleus. After treatment, the MMS was inactivated with 10% Sodium Thiosulfate and fixed with 4% formaldehyde. Cells were immobilized in agarose and 15 YFP and DAPI images were captured at 0.2-µm intervals along the z-axis with an oil immersion 100x objective on a Confocal Olympus FV1000 Upright microscope quantitated with FIJI. Images were analyzed for foci in at least three consecutive z-planes. For each sample, 300 individual cells were analyzed, and the data are represented as a percentage. The results shown are the average of three independent experiments. Merged images were created in FIJI Is Just ImageJ (FIJI) (Schindelin et al., 2012).

### Chromosome Loss Assay

An *S. cerevisiae* reporter strain containing a chromosome fragment with *SUP11* (Au et al., 2008; Hieter, Mann, Snyder, & Davis, 1985) (generously provided by Munira Basrai) plus test plasmids were grown overnight in media lacking uracil and histidine with limited adenine plus 3% raffinose. The next day, cultures were harvested and diluted down to an OD600 of 1 and then were diluted 1:1000 in water. Then 10, 30, and 50 μL of diluted cells were plated onto plates lacking uracil and containing limited adenine with either 2% glucose or 3% raffinose and 2% galactose and grown for 5 days at 30°C. Plates were then moved to 4°C for 3 days to enrich the red pigment. For each sample, 300 colonies were counted and any colony exhibiting red pigment were noted. The number of colonies with pigment was divided by the number of colonies counted. These data were generated in triplicate.

## RESULTS

### Overexpression of Ntg1 impairs *S. cerevisiae* cell growth

Our previous study demonstrates that overexpression of human NTHL1 causes early cancer phenotypes, including genomic instability, in mammalian cells (Kristin L. Limpose et al., 2018), but defining the pathways that cells employ to combat this damage is challenging in mammalian cells. Thus, we assessed whether a budding yeast model could be employed to define the pathways that contribute to phenotypes observed upon overexpression of the BER protein, NTHL1. To analyze the budding yeast counterpart of NTHL1, Ntg1 (Eide et al., 1996), we first tested whether overexpression of Ntg1 in budding yeast causes a growth phenotype. In parallel with studies of NTHL1 (Kristin L. Limpose et al., 2018), we examined overexpression of both wild-type Ntg1 and a catalytically dead variant of Ntg1 (K243Q) (Augeri et al., 1994) we term ntg1_catdead_ (D. B. Swartzlander et al., 2010; Daniel B. Swartzlander et al., 2016). We expressed these Ntg1 proteins from a galactose-inducible plasmid (Johnston, 1987) to allow regulated expression. Wild-type *S. cerevisiae* cells were transformed and grown on control glucose plates or galactose plates to induce expression of Ntg1 or ntg1_catdead_. As controls, we employed Vector alone and a subunit of the RNA exosome, Rrp44 (Schneider, Anderson, & Tollervey, 2007), which impairs cell growth when overexpressed. The Rrp44 subunit of the RNA exosome is a 3’-5’ exonuclease/endonuclease that mediates RNA decay and processing (Schneider et al., 2007; Schneider, Leung, Brown, & Tollervey, 2009). We reasoned that overexpression of this RNA processing factor impairs cell growth through mechanisms distinct from Ntg1 overexpression and thus could serve as a control for efficient galactose-mediated induction under different growth conditions.

Cells that overexpress Ntg1 show a mild impairment of growth when compared to control cells with Vector alone (Figure 1A). Cells that express ntg1_catdead_ also show an apparent slow growth (Figure 1A) compared to the Vector control cells. To provide a quantitative measure of growth defects in cells that overexpress Ntg1 or ntg1_catdead_, we conducted a liquid growth assay. As shown in Figure 1B, cells that overexpress either Ntg1 or ntg1_catdead_ show slower growth than control cells with Vector alone. Notably, overexpression of ntg1_catdead_ causes slower growth than overexpression of Ntg1 which is also reflected in the smaller colony size of ntg1_catdead_ (Figure 1A).. The difference in growth between cells overexpressing wild-type Ntg1 and ntg1_catdead_ could be explained by a difference in the level of expression achieved. To assess whether Ntg1 and ntg1_catdead_ are expressed at similar levels, wild-type cells containing a galactose inducible myc-tagged Ntg1 or ntg1_catdead_ were grown in media containing galactose, samples were collected at the indicated time points, and analyzed by immunoblotting. As shown in Figure 1C, Ntg1 and ntg1_catdead_ are expressed at approximately equal levels.

**Figure 1.**
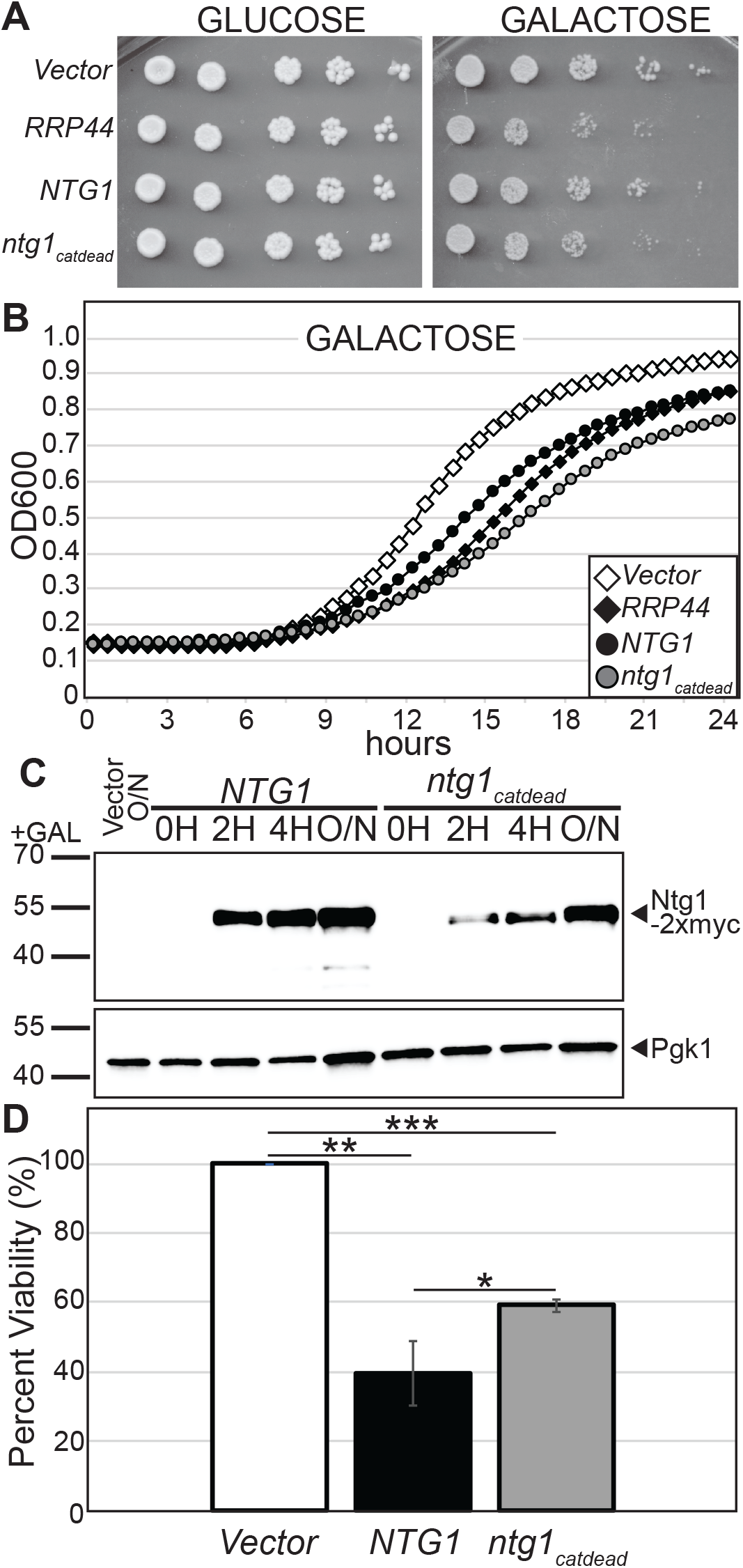
Overexpression of Ntg1 impairs *S. cerevisiae* cell growth. Wild-type cells were transformed with negative control galactose-inducible Vector (Vector), positive control galactose-inducible *RRP44*-2xmyc (*RRP44*), galactose-inducible *NTG1*-2xmyc (*NTG1*), or galactose-inducible catalytically inactive *ntg1*_*catdead*_-2xmyc (ntg1_catdead_). Cultures were grown overnight in media lacking uracil with raffinose at 30°C. A) Overnight cultures were 5-fold serially diluted and spotted on plates lacking uracil with glucose or galactose and the plates were incubated at 30°C. Pictures were taken on day 2. B) Overnight cultures were diluted into media lacking uracil with galactose and quantitative growth curve analysis was performed. OD600 readings were taken every 30 minutes and plotted vs time. Each culture appears on the graph as follows: Vector (white diamond), Rrp44 (black diamond), wild-type Ntg1 (black circle), and ntg1_catdead_ (grey circle). C) Overnight cultures were diluted into media containing galactose and samples were collected at 0, 2, 4 hours and overnight (ON). Cells were lysed and lysate was subjected to immunoblotting to detect Ntg1-2xmyc and ntg1_catdead_-2xmyc (Ntg1-2xmyc) and Pgk1 (Pgk1) serves as a loading control. Results shown in (A, B, and C) are representative of at least three independent experiments. D) Overnight cultures were diluted into media lacking uracil with galactose for 16 hours and plated on plates lacking uracil plus glucose and incubated for 4 days. At the end of 4 days, the colony forming units were counted. The viability of each sample was normalized to control Vector and is expressed as a percentage. The white circles denote the average percent viability. The * indicates a p-value of < 0.05 and ** is a p-value of < 0.005.

The growth assays cannot distinguish whether the delay in growth caused by overexpression of Ntg1 and ntg1_catdead_ is due to a cytostatic effect, where cell growth is arrested, or a cytotoxic effect, where cell viability is lost. To assess whether Ntg1 overexpression is cytostatic or cytotoxic, we conducted a viability test by inducing the expression of Ntg1 or ntg1_catdead_ with galactose overnight. After induction, cells were plated on plates containing glucose to determine the number of viable cells present in the culture. Colony forming units were counted, averaged, and compared to cells expressing the control Vector alone. The percent viability of cells expressing Vector control was set to 100%. The percent viability of colonies overexpressing Ntg1 (40%, p-value of 0.0004) or ntg1_catdead_ (59%, p-value of 0.0001), is significantly decreased when compared to Vector (Figure 1D). The viability of colonies overexpressing Ntg1 is significantly different (p-value of 0.0236) from the percent viability for cells overexpressing ntg1_catdead_. Thus, overexpression of Ntg1 and ntg1_catdead_ impair cell growth and induce cell death with a greater effect in cells overexpressing catalytically active Ntg1 as compared to a catalytically inactive variant of Ntg1.

### Overexpression of Ntg1 causes DNA double-strand breaks and chromosome loss in *S. cerevisiae*

Previous work showed that overexpression of NTHL1 causes accumulation of double-strand breaks (Kristin L. Limpose et al., 2018). To test for induction of double-strand breaks in the budding yeast model overexpressing Ntg1, we employed an *S. cerevisiae* reporter system expressing Rad52-YFP which forms foci at sites of a double-strand breaks (de Mayolo et al., 2010). We overexpressed Ntg1 or ntg1_catdead_ in these Rad52-YFP cells and analyzed the number of foci that form compared to control cells with Vector alone (Figure 2A). In Figure 2B, we quantified the percent of cells with foci. In cells with control Vector, this value was 0.11%, with cells expressing Ntg1 at 5 % (p-value of 0.016), and ntg1_catdead_ at 1.3% (p-value of 0.258). The difference in double-strand break foci detected in cells that overexpress ntg1_catdead_ is statistically different from that of foci produced in cells that overexpress wild-type Ntg1 (p-value of 0.03), consistent with a requirement for the catalytic activity of Ntg1 to cause accumulation of double-strand breaks.

**Figure 2.**
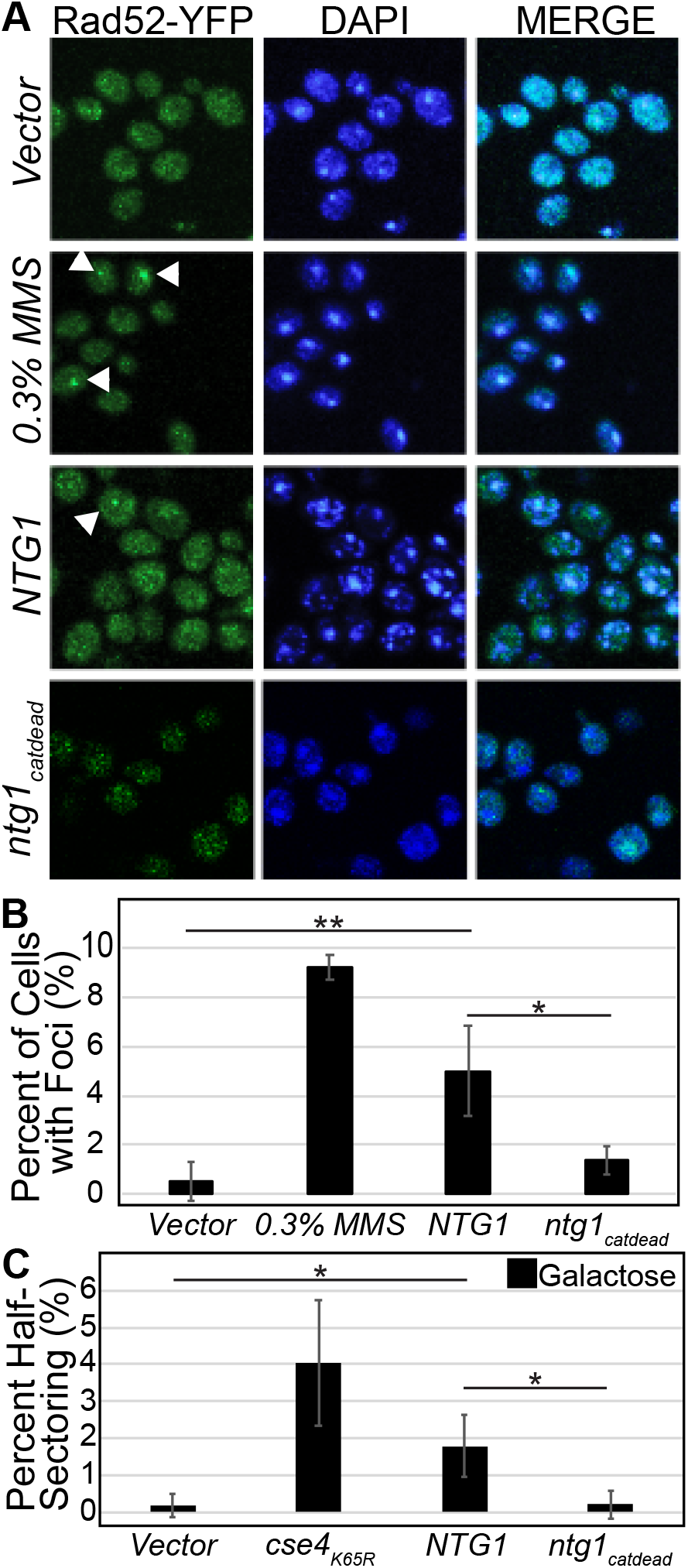
Overexpression of Ntg1 causes DNA double strand breaks and chromosome loss in *S. cerevisiae*, however, overexpression of ntg1_catdead_ does not cause chromosome loss. Cells expressing Rad52-YFP (de Mayolo et al., 2010) were transformed with negative control galactose-inducible Vector (Vector), galactose-inducible NTG1-2xmyc (NTG1), or galactose-inducible ntg1_catdead_-2xmyc (ntg1_catdead_). Cultures were grown overnight in media lacking uracil with raffinose. Samples were dosed with galactose and, as a positive control, Vector was dosed with galactose and 0.3% MMS (0.3% MMS). These cultures were incubated at 30°C for two hours. DAPI staining was utilized to visualize chromatin within the nucleus. MMS was inactivated and cells were fixed. A) Representative images from each sample show YFP (YFP), DAPI (DAPI), and merged images (MERGE). B) For each sample, 300 cells were analyzed for foci and the data are presented as the percent of cells analyzed that show foci. The results shown are the average of three independent experiments. Error bars represent standard deviation. The * indicates a p-value of < 0.05 and ** is a p-value of < 0.01.C) Cells containing a colorimetric reporter chromosome to measure chromosome loss (YPH1018 (Au et al., 2008)), were transformed with negative control galactose-inducible Vector (Vector), positive control galactose-inducible 6His-3HA-cse4K65R (*cse4K65R*), galactose-inducible NTG1-2xmyc (NTG1), or galactose-inducible ntg1_catdead_-2xmyc (ntg1catdead). Cultures were grown overnight in media lacking uracil with limited adenine and raffinose at 30°C. Cells were plated on plates lacking uracil with limited adenine and either glucose, or raffinose and galactose and the plates were incubated for 5 days at 30°C. C. C) For each sample, 300 cells were analyzed for chromosome loss and the data are represented as the percentage of cells showing chromosome loss in this assay as indicated by the number of half-sectoring colonies present in each sample, indicative of chromosome loss that occurs at the first cell division. An * indicates a p-value of < 0.05. The results shown are the average of three independent experiments.

As double-strand breaks can lead to loss of genetic material and previous analysis of overexpression of NTHL1 showed an accumulation of micronuclei (Kristin L. Limpose et al., 2018), we extended this analysis to analyze loss of whole chromosomes. To test for chromosome loss, we employed a colorimetric sectoring assay^50,55^. We overexpressed Ntg1 or ntg1_catdead_ in an *S. cerevisiae* reporter strain containing a covering chromosome fragment that when lost results in cells producing a red pigment (Au et al., 2008; Hieter et al., 1985). Figure 2C shows the percent of colonies that lost the chromosome fragment during the first division resulting in a half-sectored colony. We quantified the percent of half-sectored colonies in cells with control Vector (0.2%), Ntg1 (1.8%, p-value of 0.037) and ntg1_catdead_ (0.2%). The difference in colonies that half-sectored in cells that overexpress Ntg1 is statistically different from that of half-sectoring colonies produced in ntg1_catdead_ (p-value of 0.043). Thus, overexpression of Ntg1 but not ntg1_catdead_ results in an increase in both double-strand breaks and whole chromosome loss.

### Interplay of base excision repair with DNA damage response pathways

As we have established that overexpression of yeast Ntg1 causes a growth phenotype and exhibits similar DNA damage phenotypes as detected for overexpression of NTHL1 (Kristin L. Limpose et al., 2018), we next exploited the yeast deletion collection to interrogate pathways involved in DNA damage response for genetic interactions with overexpression of Ntg1. Because we have previously shown that human NTHL1 can potentially interact with proteins from other DNA damage response and DNA repair pathways, including nucleotide excision repair, homologous and non-homologous recombination, mutants were selected for screening that contained deletions in genes encoding proteins involved in similar yeast pathways (Limpose et al, 2017; Kar et al., 2021). In addition, since we previously reported that Ntg1 is sumoylated in response to oxidative stress, we also included genes involved in the SUMO-mediated DNA damage response (Swartzlander et al., 2016). For these studies, we employed a serial dilution and spotting assay to assess relative effects on cell growth. As described in Materials and Methods, this serial dilution assay employs 5-fold serial dilutions. This approach means that any change in growth between adjacent spots of cells on the plate reflects a five-fold change in growth. We selected a panel of deletion mutants from the yeast deletion collection and overexpressed either Ntg1 or ntg1_catdead_. We used the Vector alone as the control as well as overexpression of Rrp44. We reasoned that by setting growth in cells expressing Vector alone to a standard for each mutant analyzed, we could account for any differences in growth between wild-type cells and the deletion mutants analyzed. We employed the Rrp44 control to ensure that galactose induction is functional in each of the mutants analyzed. This approach could be employed to identify pathways that interact with overexpression of Ntg1, but also to infer which pathways might be employed for cells to respond to Ntg1-induced damage by identifying those pathways that when impaired make cells more susceptible to overexpression of Ntg1 and/or ntg1_catdead._ Such an analysis is readily performed using the budding yeast system.

Figure 3 shows examples of the serial dilution growth assays (Figure 3A,B) that were performed as well as a heat map that summarizes the complete set of deletion mutants analyzed (Figure 3C). Results of the serial dilution growth assay for a deletion mutant, *Δtel1*, that shows no genetic interaction with overexpression of either Ntg1 or ntg1_catdead_ (Figure 3A) and a deletion mutant, *Δrad51* (Figure 3B), that is sensitive to overexpression of wild-type Ntg1. Figures 3A and 3B show wild-type cells on the top panel and the deletion mutant cells on the bottom panel. The *Δtel1* cells show comparable growth to wild-type with the Vector alone control with no change in cell growth that shows any detectable difference from the wild-type control cells for either Ntg1 or ntg1_catdead_. Figure 3B shows a change observed in the growth of *Δrad51* cells that overexpress Ntg1, as evidenced by the lack of growth detected in the fourth spot in the *Δrad51* cells as compared to wild-type cells, with no detectable change for overexpression of ntg1_catdead_ compared to the wild-type control.

**Figure 3.**
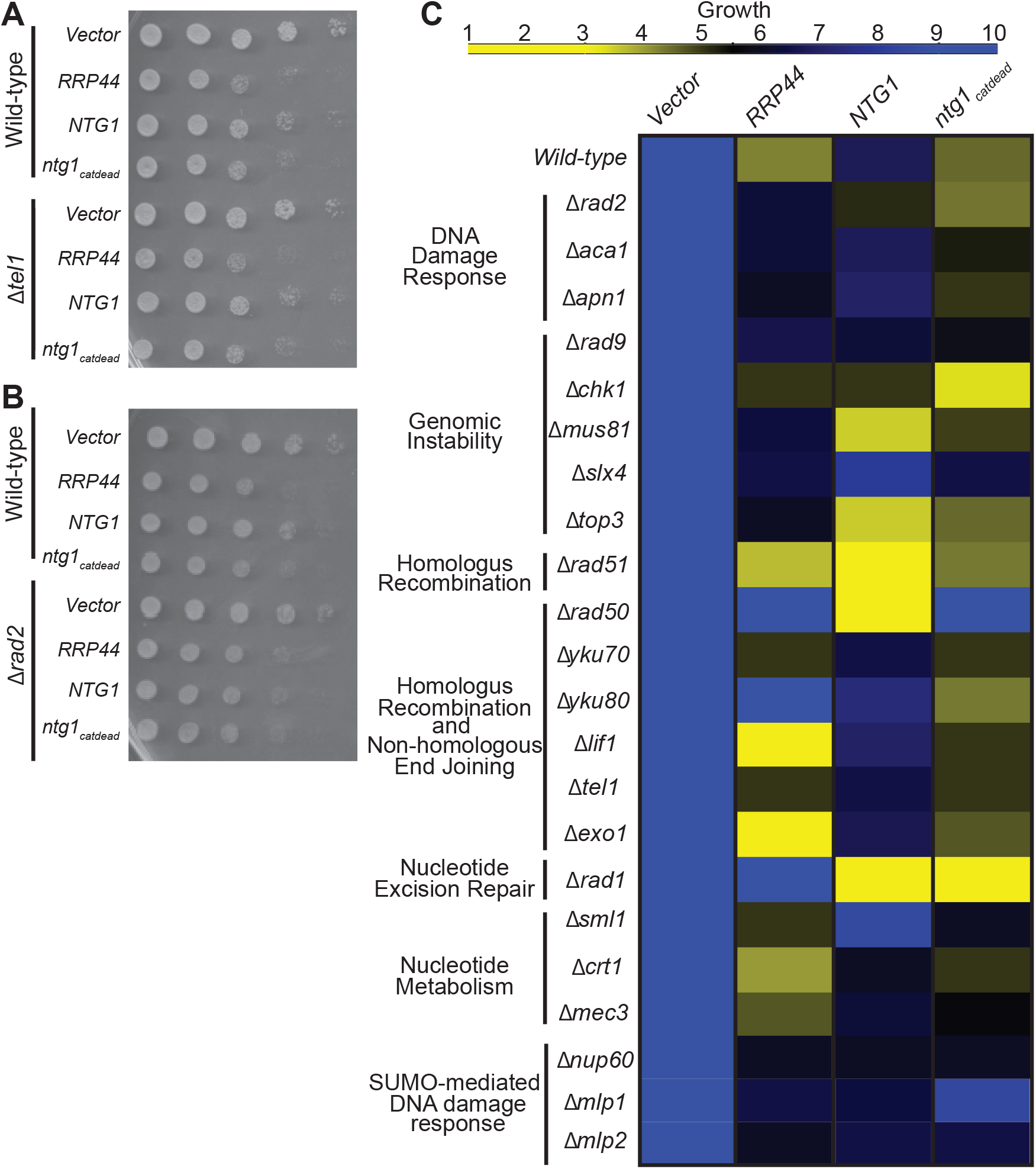
Interplay of base excision repair with DNA damage response pathways. Wild-type or a panel of S. cerevisiae deletion mutant cells were transformed with a negative control galactose-inducible Vector (Vector), positive control galactose-inducible *RRP44*-2xmyc (*RRP44*), galactose-inducible *NTG1*-2xmyc (*NTG1*), or galactose-inducible *ntg1*_*catdead*_-2xmyc (*ntg1*_*catdead*_). Cultures were grown overnight in media lacking uracil with raffinose. Overnight cultures were 5-fold serially diluted and spotted on plates lacking uracil with glucose or galactose and the plates were incubated at 30°C. Pictures were taken on day 2. Representative images from the growth assays that were employed to assess pathway interaction are shown for a mutant that (A) shows no change in growth compared to wild-type cells (*Dtel1*) and for (B) a deletion mutant (*Δrad51*) that shows increased sensitivity to overexpression of Ntg1 but not ntg1_catdead_ (*Δrad51*). C) This approach was employed to generate a heat map for a panel of deletion mutants involved in DNA damage response pathways. As described in Materials and Methods, the scale for the heat map, which is shown at the top, ranges from 1 to 10. A score of 10 means growth is detected in all five of the serially diluted spots. A score of 1 means only poor growth was detected in the most concentrated spot. “Normal growth”, which is growth detected across the serial dilution in all spots, corresponding to a score of 10, is indicated by a bright blue color on the scale. Each mutant is normalized to the respective growth of the Vector expressed in that mutant background. Only those mutants that show no change in sensitivity to overexpression of the control RNA exosome subunit, Rrp44, were considered as showing a genetic interaction with Ntg1 and/or Ntg1_catdead_. The data compiled in the heat map are representative of at least three independent experiments for each deletion mutant.

We employed these 5-fold serial dilution growth assays to develop a semi-quantitative scale, allowing us to analyze and compare results for a panel of deletion mutants. If growth was detected in all spots, this was scored a 10 on the growth scale. Growth in four spots was scored an 8, continuing to growth in a single spot set to 2. We employed a scale of 1-10 to allow for some subjective analysis of how much growth was evident in the most dilute spot where growth was detected. For all mutants analyzed, we used conditions where growth could be detected in all five spots in the Vector control sample, providing a comparable scale for all deletion mutants regardless of any growth defect present in the mutant cells. In addition, we focused our conclusions on mutants where overexpression of the galactose-inducible control, Rrp44, did not markedly change growth as compared to the wild-type cells as a control to ensure that any differences detected were not due to altered galactose-mediated induction. All results presented in Figure 3C represent the consensus result obtained from three independent growth assays.

The heat map shown in Figure 3C summarizes the data from a panel of deletion mutants analyzed. In this heat map, the growth of cells with Vector control was set to 10 (bright blue). All deletion mutants should be compared to the top row which shows the effect of overexpressing Ntg1 (7/8, dark blue), ntg1_catdead_ (5/6, gold) or the control Rrp44 (4/5, yellow gold) on this growth scale in wild-type cells. Results are shown for 22 different deletion mutants, which were divided into categories based on function. We focused on these 22 mutants to provide a representation of different DNA repair relevant pathways and illustrate how this system could be used to survey a variety of these pathways.

Based on the data summarized in Figure 3C, overexpression of both Ntg1 and ntg1_catdead_ causes slow growth comparable to that detected in wild-type cells for several of the mutants analyzed, including *Δapn1, Δyku70, Δyku80, Δtel1, Δexo1* and *Δcrt1*. Several mutants are sensitive to overexpression of both Ntg1 and ntg1_catdead_ with the most striking example the nucleotide excision repair pathway mutant, represented by *Δrad1*. We also identified pathways that are more sensitive to overexpression of wild-type Ntg1 than ntg1_catdead_. For example, both *Δrad51* and *Δrad50* cells show a stronger growth defect when Ntg1 is overexpressed as compared to wild-type cells, but no significant change as compared to wild-type cells with overexpression of ntg1_catdead_. As both Rad50 and Rad51 are key components of homologous recombination (Wright, Shah, & Heyer, 2018), this result suggests that cells with impaired homologous recombination are sensitive to overexpression of catalytically active Ntg1. Interestingly, we did not identify any mutants that show sensitivity to overexpression of ntg1_catdead_ without a concomitant effect for Ntg1. This finding is consistent with the model that overexpression of Ntg1/NTHL1 can impair cell growth and cause replication stress through at least two mechanisms, one of which results from the catalytic activity of Ntg1/NTHL1 and one that results from altered protein/protein interactions (Kristin L. Limpose et al., 2018). Catalytically active Ntg1/NTHL1 could mediate both of these effects, while ntg1_catdead_ would only impact protein/protein interactions. The broad pathways that show sensitivity to overexpression of Ntg1 and/or ntg1_catdead_ are homologous recombination and nucleotide excision repair.

Among the pathways that display some resistance to overexpression of Ntg1 and/or ntg1_catdead_, *Dmlp2* cells are more resistant to overexpression of ntg1_catdead_ than to overexpression of the control Rrp44 (Figure 3C). The Mlp2 protein is a component of the SUMO-mediated DNA damage response pathway (Strambio-de-Castillia, Blobel, & Rout, 1999), raising the possibility that SUMO modification of a protein could modulate the repair pathways by which cells respond to overexpression of Ntg1.

### SUMO modification modulates the effect of Ntg1 overexpression

A potential link to SUMO modification in cellular responses to overexpression to Ntg1 is intriguing because previous work demonstrated that Ntg1 is sumoylated, raising the possibility that sumoylation of Ntg1 could regulate Ntg1-mediated DNA damage (Daniel B. Swartzlander et al., 2016). To further investigate a potential role for sumoylation in cellular response to overexpression of Ntg1, we employed a yeast mutant lacking both SUMO E3 ligases, Siz1 and Siz2 (*Δsiz1/Δsiz2*) (Erica S. Johnson, 2004; E. S. Johnson & Gupta, 2001). The results of this analysis show that overexpression of both Ntg1 and ntg1_catdead_ in cells lacking global sumoylation causes a modest decrease in cell growth on the galactose plates relative to the wild-type control cells (Figure 4A). We also tested overexpression of Ntg1 in yeast cells with impaired desumoylase activity, by employing a temperature sensitive *ULP1* mutant (*ulp1-1*) (Li & Hochstrasser, 1999). The *ulp1-1* cells grow poorly on galactose plates; however, overexpression of Ntg1 impairs cell growth relative to Vector alone (Figure 4B). These data further suggest that SUMO-mediated interactions can contribute to the growth phenotypes caused by overexpression of Ntg1.

**Figure 4.**
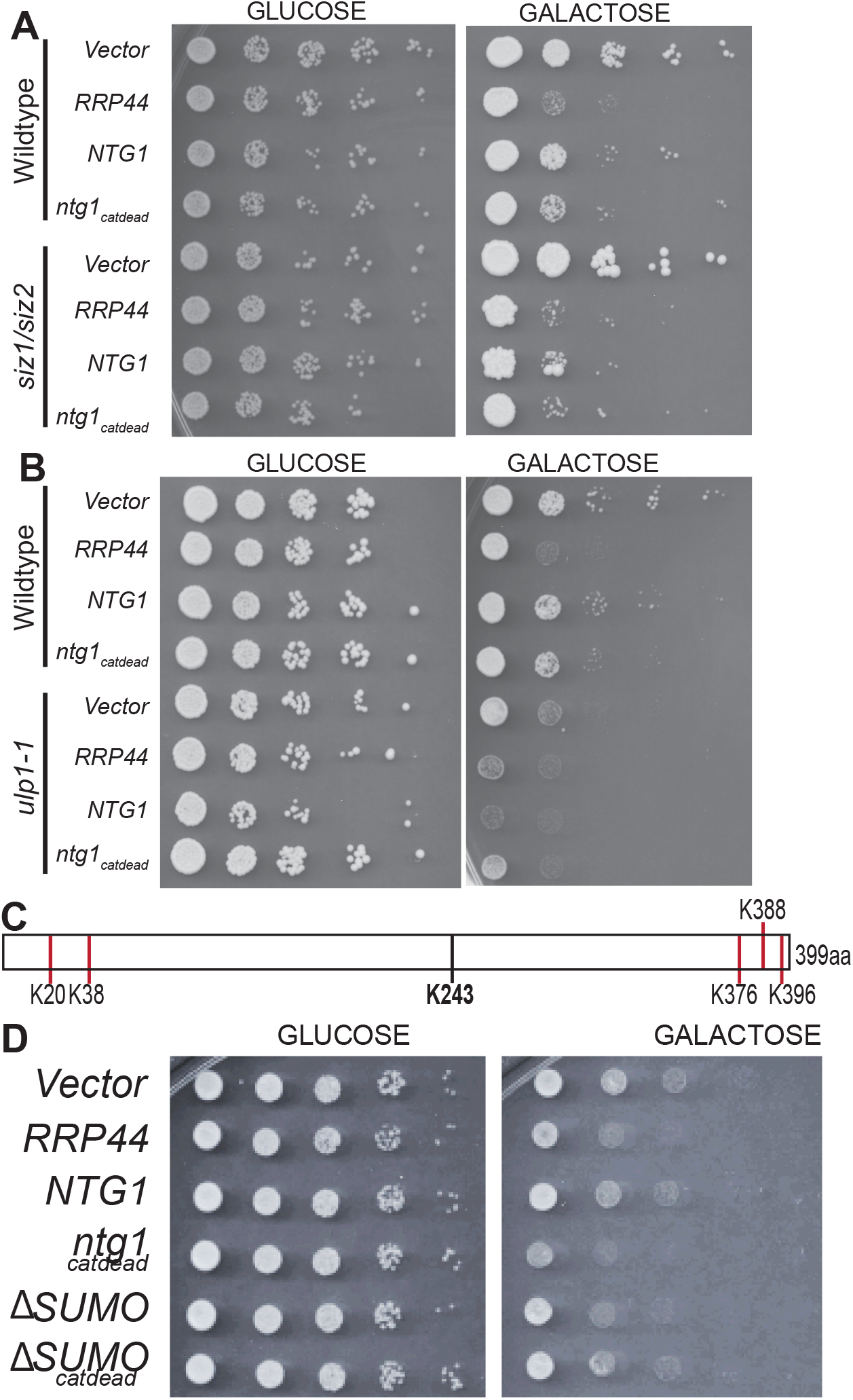
SUMO modification modulates the effect of Ntg1 overexpression. Either (A) *Δsiz1/2* or (B) *ulp1-1* cells were transformed with negative control galactose-inducible Vector (Vector), positive control galactose-inducible R*RP44*-2xmyc (*RRP44*), galactose-inducible *NTG1*-2xmyc (*NTG1*), or galactose-inducible *ntg1*_*catdead*_-2xmyc (*ntg1*_*catdead*_). Cultures were grown overnight in media lacking uracil with raffinose at 30°C. Overnight cultures were 5-fold serially diluted and spotted on plates lacking uracil with glucose or galactose and the plates were incubated at 30°C. Pictures were taken on day 2. Results shown are representative of at least three independent experiments. C) Schematic depicting the five sumoylation sites and catalytic lysine of Ntg1 (Daniel B. Swartzlander et al., 2016). The SUMO modification sites are denoted by a red line (K20, 38, 376, 388, and 396), and the catalytic lysine is indicated by a black line (K243). D) Wild-type yeast were transformed with galactose-inducible plasmids, negative control Vector (Vector), positive control *RRP44*-2xmyc (*RRP44*), *NTG1*-2xmyc (*NTG1*), ntg1_catdead_-2xmyc (*ntg1*_*catdead*_) or a nonsumoylatable ntg1 variant with or without catalytic activity, *ntg1K20,38,376,388,396R* (*ΔSUMO*), or *ntg1K20,38,376,388,396R*_*catdead*_ (*ΔSUMO*_*catdead*_). Cultures were grown overnight in media lacking uracil with raffinose at 30°C. Overnight cultures were 5-fold serially diluted and spotted on plates lacking uracil with either glucose or galactose and the plates were incubated at 30°C. Pictures were taken on day 2. Results shown in (D) are representative of at least three independent experiments.

As Ntg1 can be modified by SUMO on multiple lysine residues (Figure 4C) (Daniel B. Swartzlander et al., 2016), we directly tested whether sumoylation of Ntg1 impacts the overexpression phenotype. To address this question, we exploited a nonsumoylatable Ntg1 variant (DSUMO) where five lysines are changed to the conserved but nonsumoylatable residue arginine (K->R) (Daniel B. Swartzlander et al., 2016). Overexpression of this nonsumoylatable Ntg1 causes more of a growth defect than wild-type Ntg1 (Figure 4D). In contrast, a variant of Ntg1 that lacks SUMO modification and catalytic activity (ΔSUMO_catdead_) does not impair cell growth to the same extent as overexpression of ntg1_catdead_ (Figure 4D). These results suggest that SUMO modification of Ntg1 contributes to the overexpression phenotype of Ntg1.

## DISCUSSION

In this study, we established a budding yeast model that can be used to explore how cells respond to overexpression of Ntg1 and potentially extended to define how overexpression of NTHL1 could contribute to cancer phenotypes. We present evidence that overexpression of catalytically active Ntg1 causes an increase in both double-strand breaks and chromosome loss. In contract, while overexpression of a catalytically inactive form of Ntg1 also impairs cells growth when overexpressed, no increase in double-strand breaks or chromosome loss is evident. These results support a model where overexpression of Ntg1 can alter cell physiology through at least two mechanisms, one that depends on the enzymatic activity and one that does not. We exploited this yeast genetics system to define cellular pathways that display genetic interactions with overexpression of Ntg1. As illustrated in Figure 5, these genetic interactions identify potential interactions with DNA repair pathways that are involved in NER (Rad1), double-strand break repair (Rad50, Rad51), and SUMO-mediated DNA damage response (Mlp2), extending our previous observation that Ntg1 is sumoylated (Swartzlander et al., 2016).

**Figure 5.**
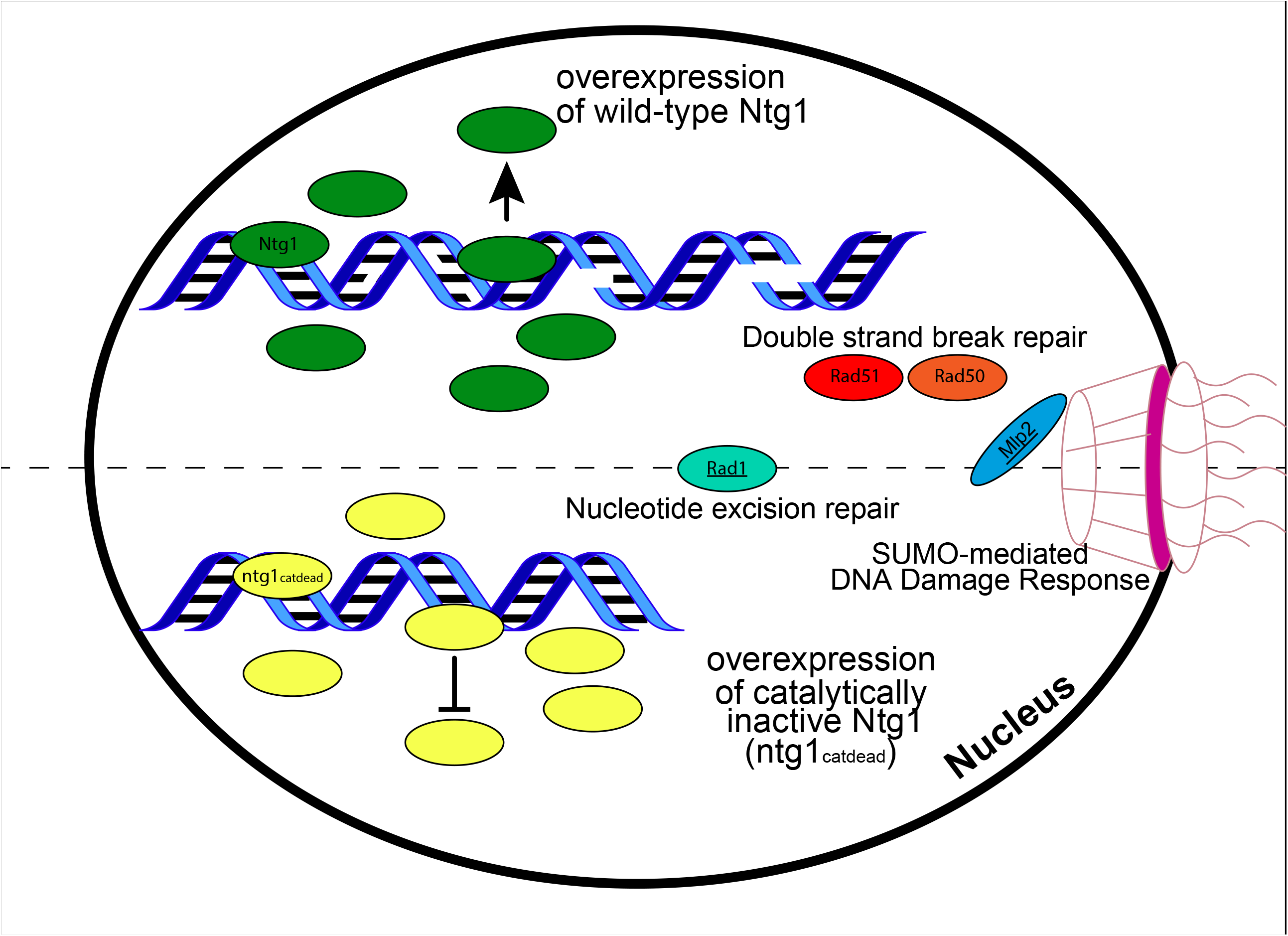
Model summarizing genetic interactions identified upon Ntg1 overexpression. A model illustrates the functional consequences of overexpression of wild-type Ntg1 (top) and catalytically inactive Ntg1, ntg1_catdead_. Cells lacking either Rad1 or Mlp2 (underlined) are hypersensitive to overexpression of both Ntg1 proteins. In contrast, only wild-type Ntg1 overexpression is toxic to cells lacking components of double-strand break repair (Rad50 or Rad51). These findings suggest that catalytically active Ntg1 has the potential to contribute to cellular phenotypes by introducing nicks into the DNA backbone that could accumulate and, ultimately, require cells to deploy homologous recombination to repair the damage. In contrast, overexpression of both wild-type and ntg1_catdead_ could lead associated proteins to be sequestered and thus decrease the capacity of other repair pathways, including nucleotide excision repair and the SUMO-dependent pathways.

The evolutionarily conserved BER proteins human NTHL1 and *S. cerevisiae* Ntg1 are functionally and mechanistically similar (Bauer et al., 2015). As described here, the similarities also extend to consequences of overexpression. Overexpression of NTHL1 results in an increase in DNA damage, DNA double-strand breaks, and micronuclei formation (Kristin L. Limpose et al., 2018). Here, utilizing budding yeast, we show that overexpression of Ntg1 in yeast also causes double strand-breaks and chromosome loss. In humans and yeast, overexpression of both NTHL1 and Ntg1 genetically interact with the double-strand break repair pathway of homologous recombination (Kristin L. Limpose et al., 2018).

As with overexpression of human NTHL1, results from this budding yeast model suggest that overexpression of Ntg1 can impact cellular physiology through at least two pathways. First, overexpression of catalytically active Ntg1 can lead to biological endpoints that likely result from accumulation of nicks in the DNA backbone. These endpoints include double-strand breaks and chromosome loss. However, a catalytically inactive Ntg1 variant (ntg1_catdead_) can also impair cell growth and previous work has demonstrated that this Ntg1 variant is not able to introduce nicks into the DNA backbone even in the presence of DNA damage (D. B. Swartzlander et al., 2010). As overexpression of Ntg1 even in the absence of catalytic activity can impair yeast cell growth, we speculate that Ntg1 can interact with other proteins to alter key protein-protein interactions. This model was also suggested for the NTHL1 protein (Kristin L. Limpose et al., 2018), building on evidence that NTHL1 physically interacts with components of the NER pathway (Chatterjee & Walker, 2017; Spivak, 2015), including XPG (Trego et al., 2016). We did not detect genetic interactions between overexpression of either wild-type Ntg1 or ntg1_catdead_ and the yeast mutant *Δrad2* (Figure 3A). *RAD2* encodes the budding yeast orthologue of XPG (O’Donovan, Davies, Moggs, West, & Wood, 1994). While this result may seem counterintuitive, if overexpression of the Ntg1 protein sequesters Rad2, cells lacking Rad2 would not be subject to any additional biological effects if Rad2 were already absent. Thus, the lack of genetic interaction with *Δrad2* remains consistent with a model where overexpression of Ntg1 could sequester components of the NER pathways, supporting the possibility of crosstalk between these two pathways (K. L. Limpose et al., 2017).

An ideal cross-species system that would take advantage of the yeast genetics approach, but employ human NTHL1, would be overexpression of NTHL1 in budding yeast. We attempted this approach, but could not drive high expression of NTHL1 in *S. cerevisiae* despite evaluating a number of promoters and plasmids. Low levels of an epitope-tagged NTHL1 could be detected when a similar galactose-inducible approach to that described here for Ntg1 was employed (data not shown). However, no effect on cell growth was detected and some mutants that are sensitive to overexpression of Ntg1 were not sensitive to the levels of NTHL1 expression that could be achieved. As an alternative approach to define pathways sensitive to overexpression of NTHL1, we have developed doxycyline-inducible, non-tumorigenic human cell lines where expression of NTHL1 can be modulated. These systems have been established both in human bronchial epithelial cells (HBEC) cells (Ramirez et al., 2004) and in a breast epithelial cell line (MCF10A) (Soule et al., 1990). A powerful approach will be to employ the yeast system described here to develop hypotheses that can be tested in these cell culture systems and eventually in mouse models.

Taken together, results from both budding yeast and human cells suggest that regulation of BER is critically important to maintain genome integrity. Either too much or too little BER activity can be detrimental to cells, demonstrating that cells require a carefully regulated level of BER activity (Hassall, 1904).

## ACKNOWLEDGMENTS

This study was supported in part by the Emory Integrated Genomics Core (EIGC), which is subsidized by the Emory University School of Medicine and is one of the Emory Integrated Core Facilities. Additional support was provided by the Georgia Clinical & Translational Science Alliance of the National Institutes of Health under Award Number UL1TR002378. The content is solely the responsibility of the authors and does not necessarily reflect the official views of the National Institutes of Health. This research project was supported in part by the Emory University Integrated Cellular Imaging Microscopy Core. ZJ was supported by the Emory IMSD grant (5R25GM125598). AMD was supported by NIH F31 (GM115178). PWD was supported by US National Institute of Health Intramural Research Program Project Z1AES103328.

